# Inferring the infant pain experience: a translational fMRI-based signature study

**DOI:** 10.1101/2020.04.01.998864

**Authors:** Eugene P. Duff, Fiona Moultrie, Marianne van der Vaart, Sezgi Goksan, Alexandra Abos, Sean P. Fitzgibbon, Luke Baxter, Tor D. Wager, Rebeccah Slater

## Abstract

**Background:** In the absence of verbal communication it is challenging to infer an individual’s sensory and emotional experience. In adults, fMRI has been used to develop multivariate brain activity signatures, which reliably capture elements of human pain experience. We translate whole-brain fMRI signatures that encode pain perception in adults to the newborn infant brain, to advance understanding of functional brain development and pain perception in early life.

**Methods:** A cohort of adults (n=10; mean age=28.3 years) and 2 cohorts of healthy infants (Cohort A: n=15; Cohort B: n=22; mean postnatal age=3 days) were stimulated with low intensity nociceptive stimuli (64-512mN) during acquisition of functional MRI data. fMRI pain signatures were applied directly to the adult data and transformed such that they could be applied to the infant brain. In each cohort, we assessed the concordance of the signatures with the brain responses using cosine-similarity scores, and we assessed stimulus intensity encoding of the signature responses using Spearman rank correlation. Brain activity in ‘pro-pain’ and ‘anti-pain’ brain regions were also examined.

**Findings:** The Neurologic Pain Signature (NPS), which reflects aspects of nociceptive pain experience, was activated in both the adults and infants, and reliably encoded stimulus intensity. However, the Stimulus Intensity Independent Pain Signature (SIIPS1), which reflects higher-level cognitive modulation of nociceptive input, was only expressed in adults. ‘Pro-pain’ brain regions showed similar activation patterns in adults and infants, whereas, ‘anti-pain’ brain regions exhibited divergent responses.

**Interpretation:** Basic intensity encoding of nociceptive information is similar in adults and infants. However, translation of adult brain signatures into infants reveals significant differences in infant cerebral processing of nociceptive information, which may reflect their lack of expectation, motivation and contextualisation. This study expands the use of brain activity pain signatures to non-verbal patients and provides a potential approach to assess analgesic interventions in infancy.

**Funding:** This work was funded by Wellcome (Senior Research Fellowship awarded to Prof. Rebeccah Slater) and SSNAP “Support for Sick and Newborn Infants and their Parents” Medical Research Fund (University of Oxford Excellence Fellowship awarded to Dr Eugene Duff).

**Research in context:** *Evidence before this study:* We searched PubMed for research articles published prior to March 2020 using terms including ‘fMRI’, ‘infant or neonate’, and ‘pain or nociception’ in the title or abstract. Due to the relatively new emergence of this field, and the experimental and analytical challenges involved in studying cerebral processing of pain in the MRI environment in healthy newborn infants, only five fMRI studies have examined infant brain responses to nociceptive input. In a foundational pilot study, Williams et al., applied an experimental noxious stimulus to a single infant, evoking widespread brain activity that included several brain regions involved in pain processing in adults. Goksan et al., subsequently performed an observational cohort study and used regional analyses to compare active brain regions in infants (n=10) and adults (n=10), concluding that the evoked patterns of brain activity were broadly similar in infants and adults. Further follow-up analysis in the infant cohort revealed that the functional connectivity of brain regions involved in descending pain modulation influences the magnitude of pain-related brain activity. Two further studies focused on methodological advances, providing evidence-based recommendations for fMRI acquisition parameters and image processing in order to maximise the quality of infant data, and these methods have been implemented in this study.

*Added value of this study:* This study translates validated adult pain fMRI brain signatures to a nonverbal patient population in which the assessment and management of pain presents a significant clinical challenge. Application of fMRI brain signatures to newborn infants expands on previous fMRI studies that provided only qualitative evidence that noxious stimulation commonly activates brain regions in the adult and infant brain. Here we demonstrate that the basic encoding of the sensory discriminative aspects of pain, as represented by the Neurologic Pain Signature (NPS), occurs in both adults and infants, whereas higher-level cognitive modulation of pain, represented by the Stimulus Intensity Independent Pain Signature (SIIPS1) is only present in adults and not observed in infants. The differences in how the immature infant brain processes pain, relative to the mature adult brain, are likely to reflect differences in their expectation, motivation and contextualisation of external events rather than differences in their core nociceptive cerebral processing of pain. This work allows us to use quantitative fMRI observations to make stronger inferences related to pain experience in nonverbal infants.

*Implications of all the available evidence:* Behavioural pain scores used in neonatal clinical care offer limited sensitivity and specificity to pain. Neonatal clinical trials that use these scores as outcome measures frequently report a lack of efficacy of common analgesic interventions, resulting in few evidence-based drugs for treating pain. The value of using brain-based neuroimaging markers of pain as a means of providing objective evidence of analgesic efficacy in early proof of concept studies is well recognised in adults, even in the absence of behavioural pain modulation. Similarly, in infants EEG-based measures of noxious-evoked brain activity have been used as outcome measures in clinical trials of analgesics to overcome some of the inherent limitations of using behavioural observations to quantify analgesic efficacy. Considering the successful translation of the Neurologic Pain Signature (NPS) and its sensitivity to analgesic modulation in adults, this novel methodology represents an objective brain-based fMRI approach that could be used to advance the discovery and assessment of analgesic interventions in infancy.

## Introduction

In the absence of verbal communication, the perceptions of infants are enigmatic. Behavioural cues such as grimacing and vocalisation are relied upon by caregivers and clinicians to make inferences regarding an infant’s experience of pain^1^, which are broadly based on experiences of older children and adults capable of verbal report. Nevertheless, behaviours have limited sensitivity and specificity and their interpretation is subjective. Behavioural responses attributed to pain can be elicited by non-noxious stimuli in premature infants^2^ and infants can have high behavioural pain scores in response to innocuous procedures such as the changing of a nappy^3^. While pain scoring systems such as the Premature Infant Pain Profile-Revised (PIPP-R) have some construct validity^1^, the extent to which these scores correspond to pain perception is unknown. Given that pain perception is encoded by the brain, neuroimaging provides a proxy measure of neural activity that can be compared between infants and adults in order to make valuable inferences regarding the experience of pain in this non-verbal population.

Pain has no specific nexus in the human brain. Instead, a disperse mode of neural activity across sensory, limbic and other areas, with complex temporal and stochastic characteristics is thought to underlie an individual’s subjective perception^4^. The adult experience of pain involves complex cognitive and emotional processes, which are influenced by anticipation, attention, memory, salience, attribution of causality, fear and anxiety^5^. However, many of these modulatory factors may not influence pain perception in early life. As the complexity of brain circuitry develops in terms of both structural and functional connectivity, a richer experience of pain will likely evolve. Functional magnetic resonance imaging (fMRI) has been extensively used to characterise the nociceptive system in adults, and in infants noxious stimulus-evoked neural activity can be reliably detected^6,7^. Regional analysis of adult and infant responses has previously revealed broadly similar patterns of noxious-evoked activity^6^, with infants displaying more bilateral activation, which may be due to exuberant cortico-cortical and interhemispheric connectivity of the immature brain. Additional brain regions were also activated in infants, such as the auditory cortices, hippocampus and caudate, and no activity was reported in some brain regions such as the amygdala and orbitofrontal cortex. However, inferring the perceptual consequences of functional activity or inactivity within individual anatomical regions remains challenging.

Regionalised assessments of activity can be combined to make more robust inferences from data. Multivariate spatial patterns of fMRI activity have been identified that can predict specific aspects of a reported experience^8^. Recently, a series of studies in adults have characterised fMRI response signatures associated with pain and negative affect^9–14^. The primary pain template was the Neurologic Pain signature (NPS), which predicts reported pain intensity^12^. The NPS was trained on responses to varying levels of heat pain and has subsequently been shown to predict variations in pain in over 40 independent published cohorts^8^ including noxious electrical and mechanical^11^ stimuli. This signature is sensitive to nociceptive pain and does not correspond to activity evoked by non-noxious control stimuli^12^, vicarious pain^11^, threat^11^, negative affect^9^ or social pain^12^. A second signature, the Stimulus Intensity Independent Pain Signature (SIIPS1), was subsequently developed to explain additional variation in reported pain beyond the NPS and stimulus intensity, relating to modulatory factors such as expectation, motivation, context and perceived control^13^. Whether noxious-evoked brain activity in infants reflects these signatures of pain is not known.

Verbal report is the gold standard measure of pain in adults, but as infants cannot use language to describe their pain, there is great value in identifying alternative objective surrogate measures that can be used to quantify pain experience in infants. In this study, we applied graded acute mechanical nociceptive stimuli to infants and adults during fMRI acquisition and projected functional neuroimaging signatures of pain onto evoked brain activity to detect and quantify neural patterns of activity underpinning aspects of pain experience. We investigated the concordance of pain-related signatures (NPS and SIIPS1) with noxious-evoked brain activity in adults and infants, compared their expression across stimulus intensities, and examined the responses within ‘pro-pain’ and ‘anti-pain’ components of the signatures.

## Methods

### Patient cohorts and experimental protocol

This study included a cohort of healthy adults and two cohorts of healthy newborn infants scanned within 10 days of birth. Adult participants were recruited from the University of Oxford and infants from the postnatal wards of the John Radcliffe Hospital, Oxford, UK. Informed written consent was provided by adult participants and by the parents of infants prior to study commencement. All data were acquired at the FMRIB Centre, University of Oxford. The infants were studied under ethical approval from the National Research Ethics Service and the adult study was granted approval by the University of Oxford Central University Research Ethics Committee. Both studies were conducted in accordance with the Declaration of Helsinki and Good Clinical Practice guidelines.

Adult participants and infants in Cohort A were scanned in a 3T Siemens Verio scanner. The second infant cohort (Cohort B) was scanned in a newly installed Siemens Prisma 3T scanner. All infants were wrapped, fed and fitted with appropriate ear protection prior to scanning. During image acquisition, acute noxious experimental punctate stimuli of various intensities were applied ten times to the left foot of each participant (infants: 64 and 128 mN; adults: 64, 128, 256 and 512 mN; Fig. 1A) by a trained experimenter with a minimum inter-stimulus interval of 25s (Fig. 1A). The order of scans was randomised across subjects and stimuli were time-locked to the fMRI recording using Neurobehavioural Systems (Presentation, https://www.neurobs.com) software. After the scanning session, the stimuli were re-applied to the adults outside the scanner environment in order to record verbal pain scores and complete a McGill Pain Questionnaire.

**Fig. 1.**
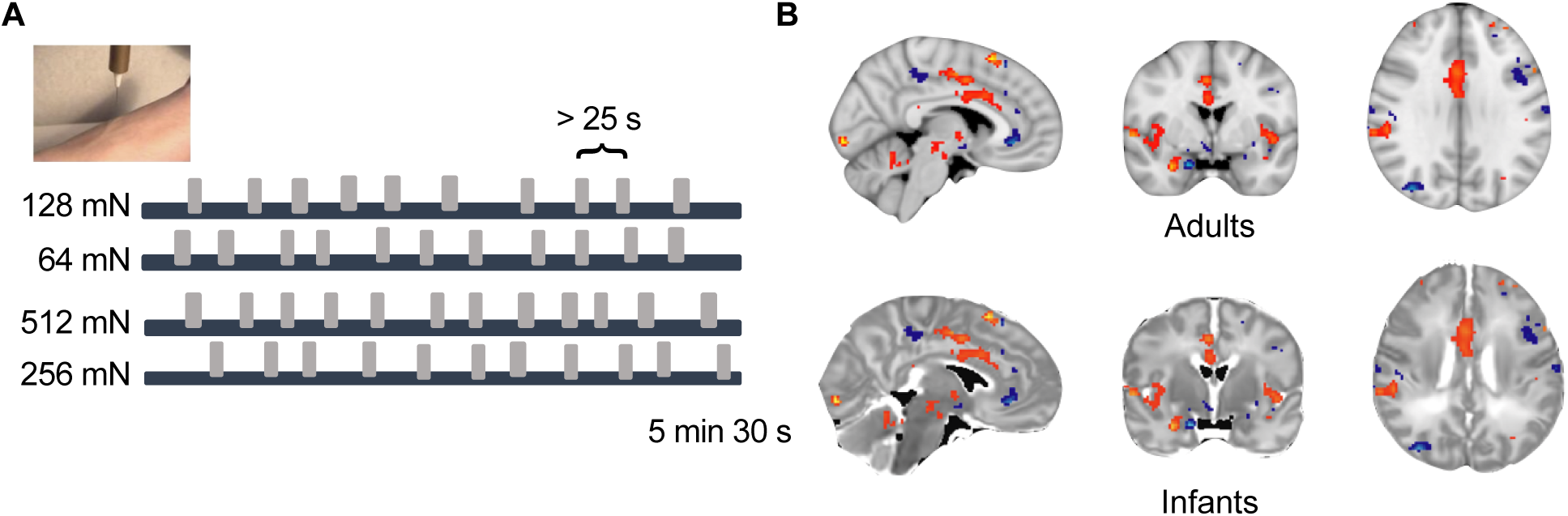
**A**. Experimental design: stimuli were applied with a minimum inter-stimulus interval of 25s across a range of intensities in a randomised order. Four stimulus intensities were used in adults and the lowest 2 stimulus intensities were used in infants. **B**. Registration of the NPS in the infant and adult. The NPS is overlaid on the registered adult and infant templates, with both templates transformed into MNI template space (see Supplementary Material for transformation methodology).

### fMRI pre-processing

The adult data were processed using FSL 6.0^15^. BOLD data were motion and distortion corrected, high pass temporally filtered with a cut-off of 100s, 5mm spatial smoothing was applied, an ICA-based clean-up was employed, and motion parameters were regressed from the data. All infant fMRI data were processed using the dHCP functional pipeline^16,17^ (see Supplementary Material) and 3mm spatial smoothing and 100s high-pass temporal filtering were applied. The pain responses were modelled using a general linear model (GLM) in FEAT^15^. In adults, the experimental design was convolved with the canonical adult double gamma HRF, and in infants the experimental design was convolved with term-infant-specific optimal basis functions generated by Arichi and colleagues^18^. Parameter maps were generated for each adult and infant at each stimulus intensity and transformed into standard Montreal Neurological Institute (MNI) template space via a series of transformations (Figure 1B; see Supplementary Material).

### Signature Analysis

In the adult cohort, six brain signatures were first applied to the fMRI response data, including the Neurologic Pain Signature (NPS)^12^ and the Stimulus Intensity Independent Pain Signature (SIIPS1)^13^, and four control signatures, which comprised of signatures of vicarious pain (VPS)^11^, picture induced negative affect (PINES)^9^, social rejection (SR)^14^ and a signature representing global signal (GS). We analysed the correspondence of each signature to the parameter maps of each stimulus intensity using a modified pipeline from the CANlab toolbox (https://canlab.github.io). Concordance between brain responses and signatures was defined by cosine-similarity scores (range: 0-1) between the parameter maps and the predefined spatial signature images.

In this study, adult data were used to validate the signature responses for our specific experimental paradigm involving application of mechanical noxious stimuli, to identify responsive subregions for infant analyses, and to facilitate direct comparison of cosine similarities between adult and infant responses. Only signatures with significant concordance in adults were applied to the infant data. For each signature, Spearman rank correlation across stimulus levels was also used to assess intensity encoding of the responses.

The NPS extends across numerous brain regions and there is spatial variation in the strength of expression of the signature^12,13,19^. Consequently, the signature has been divided into a ‘pro-pain’ component where the signature has positive weights (indicating a positive relationship with pain report) and an ‘anti-pain’ component where the signature has negative weights (indicating a negative relationship with pain report)^19^. In the NPS, the ‘pro-pain’ brain regions include the insular, secondary somatosensory, and anterior cingulate cortices (ACC), and the ‘anti-pain’ brain regions include the inferior parietal lobule and lateral occipital cortex^19^. SIIPS1 has also been divided into a ‘pro-pain’ (including the insular cortex and central operculum), and an ‘anti-pain’ component (including areas of the somatomotor cortex, hippocampus and cuneus)^13^. Therefore, secondary analyses in the adult and infant cohorts were performed examining the concordance of ‘pro-pain’ and ‘anti-pain’ subcomponents of each signature. These subcomponents have previously been used in studies of the NPS^12^ and SIIPS1^13^ and were obtained using the Canlab toolbox. Brain regions with negligible or negative signature responses to the nociceptive stimulus in the adult dataset (average cosine similarity < 0.01) were excluded from analyses.

## Results

Data from 10 healthy adults (mean age = 28 [range: 23–36] years; number of females: 5) and two cohorts of healthy newborn infants were included in this study. There were 15 infants in ‘Cohort A’ (mean gestational age (GA) at birth = 39.7 [range: 35-42] weeks; mean postnatal age (PNA) = 4 days [range:1-11]); number of females: 6) and 22 infants in ‘Cohort B’ (mean GA at birth = 39.0 [range: 36-42] weeks; mean PNA: 3 days [range: 1-10]).

Adults reported that the punctate stimuli evoked a low to moderate pain sensation, which they most frequently described as pricking (n = 8 of 10) and sharp (n = 6 of 10), and verbal pain scores increased with stimulus intensity (Spearman r=0.89; p < 0.001; Fig. 2A)^6^. The NPS and SIIPS1 signatures showed robust expression in the adults. However, the four control signatures (VPS, PINES, SR and GS) were not significantly expressed, confirming that the brain activity evoked by the physical noxious punctate stimulus most closely resembled the neurologic and stimulus independent pain signatures (Fig. 2B). The NPS responses tracked the mean reported pain intensity scores in adults (r = 0.61, p<0.001), while the SIIPS1 did not show significant association with verbal pain report (r=0.14, p=0.37).

**Fig. 2.**
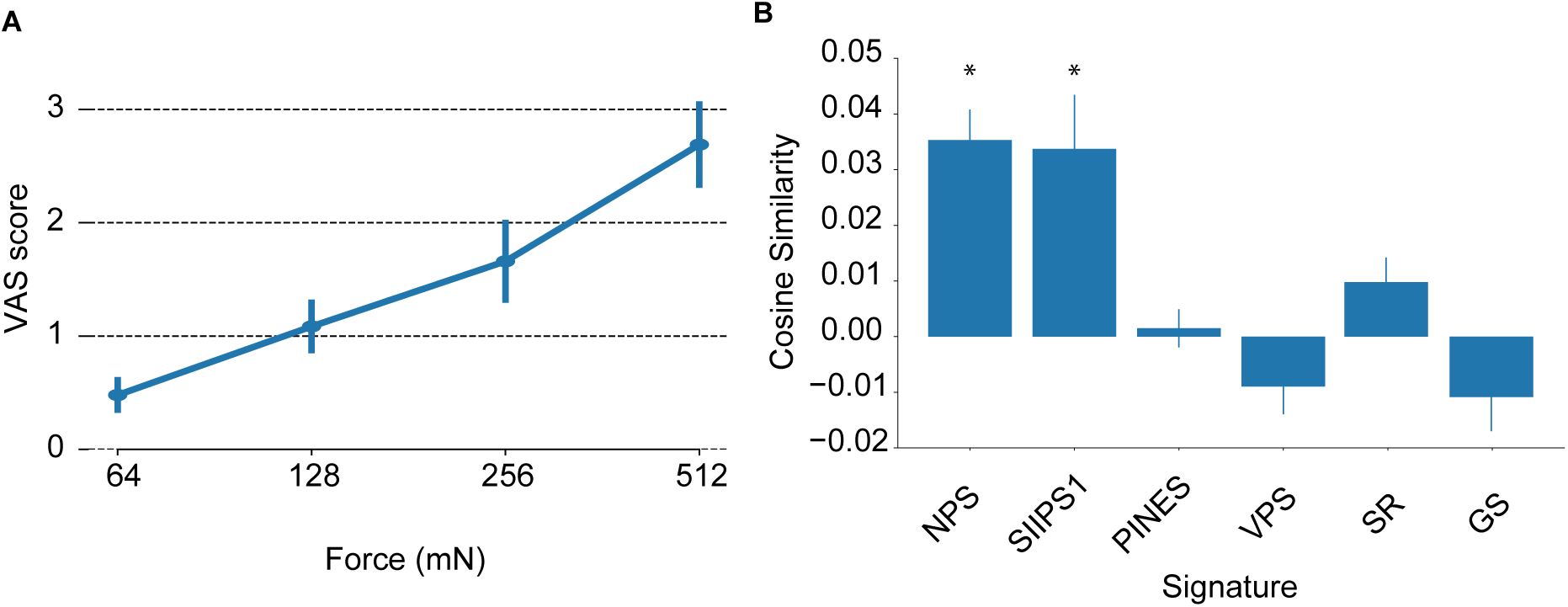
Adult Cohort **A**. Out-of-scanner pain ratings. Intensity of the verbal pain report (Visual Analogue Scale (VAS)) significantly increased with increasing stimulus intensity (r=0.89; p < 0.001). **B**. Quantification of the presence of six signatures in response to nociceptive input. Only the NPS and SIIPS1 show significant expression. * indicates cosine similarity significantly greater than zero. Abbreviations: Neurologic Pain Signature (NPS); Stimulus Intensity Independent Pain Signature (SIIPS1); and four control signatures: picture induced negative affect (PINES); vicarious pain (VPS); social rejection (SR) and global signal (GS).

We considered the comparative expression of the NPS in the adult and infant cohorts. The NPS was consistently expressed in adults (mean cosine similarity range: 0.01-0.06), and showed significant intensity encoding between 64 and 512 mN (p<0.001, Fig. 3A), consistent with its correlation with verbal pain report (Fig. 2A). Similarly, in both infant cohorts, application of the 128 mN stimuli to the foot evoked a change in brain activity that had strong concordance with the NPS (Cohort A: p=0.001; Cohort B: p=0.048; Fig. 3A). Mean cosine similarity levels in infants (at both 64 mN and 128 mN) were similar to adult values (mean cosine similarity range: 0.01-0.02). In infants, the responses following the 128 mN stimuli were greater than the 64 mN stimuli, although this was not significantly different. (Fig. 3A)

**Fig. 3.**
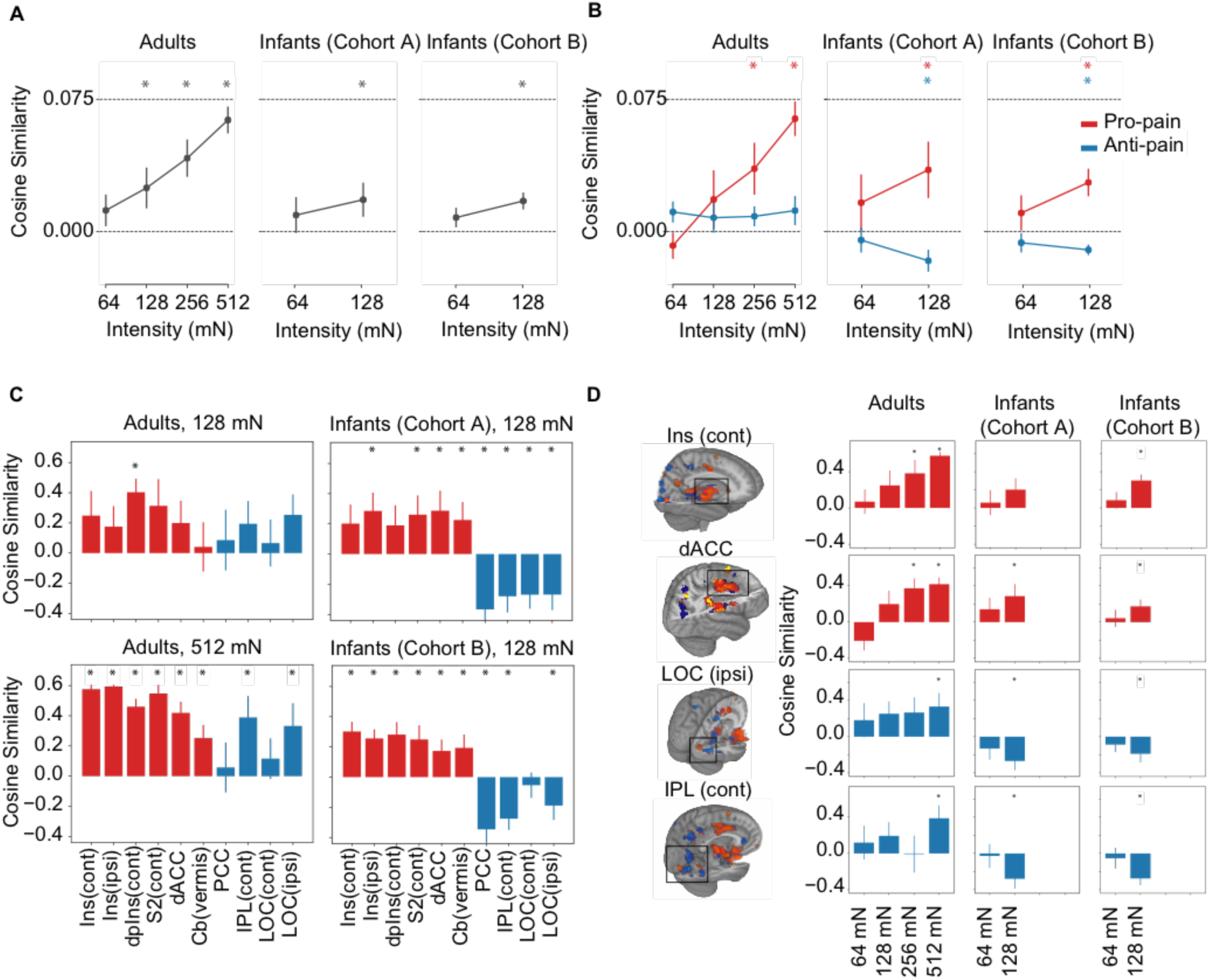
**A**. Neurologic Pain Signature (NPS) responses (measured by cosine similarity) in the Adult Cohort and two Infant Cohorts. All three cohorts showed positive cosine similarities with evidence of intensity encoding. **B**. NPS separated into ‘pro-pain’ and ‘anti-pain’ components. **C**. NPS subregion analysis. Mean NPS responses across different anatomical subregions following application of the 128 mN force (adult and infant cohorts) and 512 mN force (adult cohort only). Adults and infants show concordance with the NPS in ‘pro-pain’ subregions (red); only the adults show concordance with the NPS in the ‘anti-pain’ subregions (blue) **D**. Example intensity encoding of the NPS in selected subregions. Abbreviations: ipsilateral (ipsi); contralateral (cont); insula (ins); dorsal posterior insula (dpIns); secondary somatosensory cortex (S2); dorsal anterior cingulate cortex (dACC); cerebellum (Cb); posterior cingulate cortex (PCC); inferior parietal lobule (IPL); lateral occipital cortex (LOC). * indicates cosine similarity significantly different from zero.

In the ‘pro-pain’ NPS component, both adults and infants showed strong intensity-dependent activation (Fig. 3B, red lines), with infants displaying greater concordance with the signature compared to adults in response to the same intensity stimulation. In adults, individual ‘pro-pain’ anatomical subregion analysis also showed positive responses, which were consistently observed in both infant cohorts (Fig. 3C, red bars). In general, activity increased with stimulus intensity within these brain regions in both the adult and infant cohorts (Fig. 3D, red bars).

The ‘anti-pain’ component of the NPS represents brain regions where an increase in pain report is associated with a decrease in brain activity. In the ‘anti-pain’ component the evoked brain activity in adults was concordant with the NPS but comparatively weaker as compared with the ‘pro-pain’ component (Fig. 3B adults, blue line). However, many individual ‘anti-pain’ brain regions, including the posterior cingulate cortex, the inferior parietal lobule and the lateral occipital cortex, showed significant expression of the NPS in response to the noxious stimuli (Fig. 3C, 3D adults – blue bars). In contrast, the infant cohorts displayed negligible cosine similarity at 64 mN intensity, and negative cosine similarity at 128 mN (Figure 3B, blue line). Regional analysis of the ‘anti-pain’ brain regions that were significantly concordant with the NPS in adults revealed a lack of concordance in infants (Fig. 3C, 3D infants – blue bars). Conversely, an inverted response relative to the NPS template was observed (Figure 3C infants – blue bars), corresponding to an increase in activation of these brain regions in response to the noxious stimulation.

The SIIPS1 signature tracks variability in subjective pain report that is not accounted for by stimulus intensity and the NPS, which tracks verbally reported pain intensity. SIIPS was reliably expressed in adults (cosine similarity: 0.023-0.062) and as expected did not show significant intensity encoding (Fig. 4A, adults). In contrast, noxious-evoked brain activity in infants was not concordant with the SIIPS1 signature (Fig. 4A, infants) and infants in Cohort B displayed a negative concordance with the signature when stimulated with the 128 mN force (Figure 4A, infants – Cohort B). However, in the ‘pro-pain’ subdivision of the signature, both adults and infants showed a similar pattern of SIIPS1 related activation (Fig. 4B, red lines) and subregion decomposition of the SIIPS1 ‘pro-pain’ component revealed that brain regions reflecting the signature in adults showed similar responses in infants (Fig. 4C, red bars). This suggests that the lack of representation of the overall SIIPS signature in the infants is primarily driven by a lack of concordance with the signature in the ‘anti-pain’ brain regions. In line with this interpretation, negative concordance with the SIIPS1 was observed in both infant cohorts when a force of 128 mN was applied, which was opposite to that which was observed in the adults (Fig. 4C). This divergence of adult and infant responses to the nociceptive input was observed across all subregions (Fig. 4C and Fig. 4D).

**Fig. 4.**
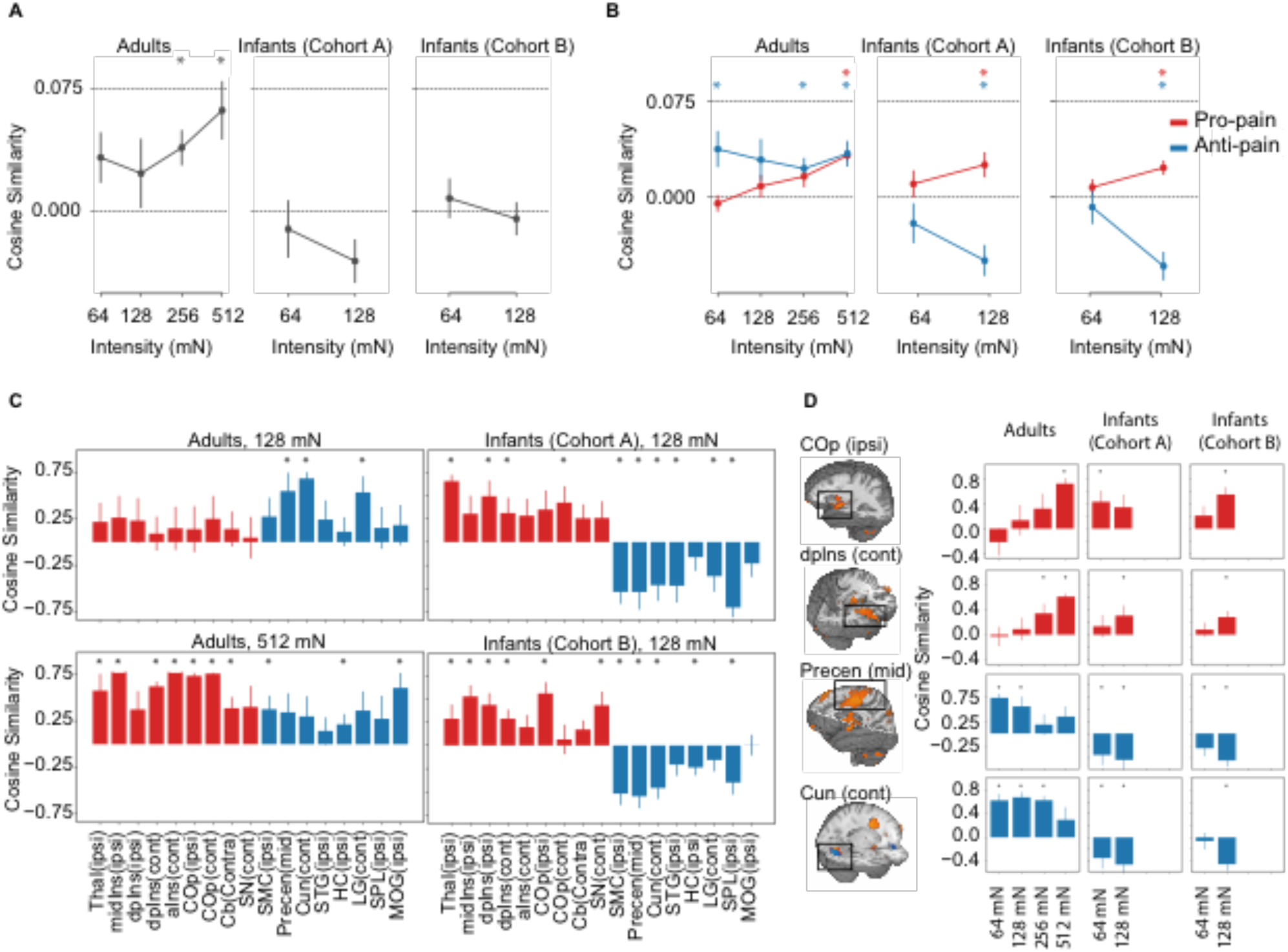
**A**. SIIPS1 responses (measured by cosine similarity) in the Adult Cohort and two Infant Cohorts. SIIPS1 signatures responses across stimulus intensity levels for the adult cohort and two infant cohorts. The adult cohort showed positive concordance with the SIIPS1 whereas the infant cohorts did not. **B**. SIIPS1 separated into “pro-pain” and “anti-pain” components. **C**. SIIPS1 subregion analysis. Mean SIIPS1 responses across different anatomical subregions following application of the 128 mN force (adult and infant cohorts) and 512 mN force (adult cohort only). Adults and infants show concordance with the SIIPS in ‘pro-pain’ subregions (red); only the adults show concordance with the SIIPS in the ‘anti-pain’ subregions (blue). **D**. Example intensity encoding of the SIIPS1 in selected subregions. Adults show evidence of intensity encoding in ‘pro-pain’ subregions. Abbreviations: ipsilateral (ipsi); contralateral (cont); thalamus (Thal); middle insula (midIns); dorsal posterior insula (dpIns); anterior insula (aINS); central operculum (COp); cerebellum (Cb); substantia nigra (SN); somatomotor cortex (SMC); precentral gyrus (Precen); Cuneus (Cun); superior temporal gyrus (STG); hippocampus (HC); lingual gyrus (LG); superior parietal lobule (SPL); middle occipital gyrus (MOG). * indicates cosine similarity significantly different from zero.

## Discussion

Multivariate fMRI signatures have been designed to robustly link features of brain activity to subjective experiences^8,12^, and a fundamental potential use of these signatures is their application to challenging patient populations with limited communication. In this study, we translated whole-brain fMRI signatures that predict aspects of pain perception in adults to the newborn infant brain, in order to advance our understanding of how infants experience noxious events in early life. Data from our adult cohort first confirmed that brain activity evoked by the brief mechanical nociceptive stimulus applied in this study was significantly concordant with two validated brain signatures of pain, the NPS and SIIPS1. However, there was no concordance of evoked brain activity with control signatures of vicarious pain, negative affect or global brain signals, providing further evidence of the specificity of these signatures. In two independent cohorts of infants, the NPS, a signature which tracks reported pain intensity across a variety of nociceptive stimuli in adults^11,12^, was significantly expressed in infant evoked brain activity. However, the SIIPS1 signature, which quantifies additional cerebral contributions to reported pain experience beyond nociceptive input and intensity encoding^13^, was not observed in infants. Furthermore, ‘pro-pain’ NPS brain regions, primarily targeted by ascending nociceptive afferents, showed similar activation patterns in adults and infants, whereas, ‘anti-pain’ regions, indirectly targeted by nociceptive afferents, showed divergent responses between the adult and infant. Overall, these results suggest that the neural activity patterns that are indicative of intensity-modulated pain in adults, evidenced by correlation with verbal report, are also present in infants.

Human language provides an exquisite means by which subtle sensory and affective states can be communicated. However, in non-verbal patient populations, such as newborn infants, inference about sensory and emotional experiences are guided by behavioural and physiological observations. This approach is relied upon in clinical practice for the quantification and treatment of infant pain, but these measures are subjectively recorded and lack specificity. The relationship between observed behaviours and conscious experience of pain is ultimately unknown. Neuroimaging provides more direct insight into the neural processes underlying infant experience and has the potential to improve the inferences that can be made regarding pain and analgesia in nonverbal populations. In infants, noxious-evoked behaviour and brain activity have been studied and correlated^20–22^. However, determining a quantitative association between behaviour, or brain activity, and perception has remained elusive. Unlike previous infant fMRI studies that have focused on regional analyses of individual brain regions associated with pain in adults^6^, multivariate fMRI signatures incorporating combinations of regional activity allow for more accurate and specific prediction of pain reports^8,13^. Our results provide quantitative evidence that noxious stimulation in infants evokes changes in brain activity that would be attributed to the sensory discriminative aspects of pain experience in adults. The similar expression of the NPS in both adults and infants, coupled with the differences in the SIIPS expression across these cohorts, is consistent with the results of a recent study comparing adult and infant pain responses using EEG^20^. In this study, infants displayed noxious evoked potentials and gamma oscillations, which are also markers of primary nociceptive processing and subjective pain in adults^23^.

In contrast, nociceptive stimulation in newborn infants did not evoke patterns of cerebral activity characterised by the SIIPS1, which relate to complex psychological constructs in adults, including expectation, motivation, contextualisation and perceived control of pain^13^. During early development, infants become gradually more engaged with their external environment but it is not known at what developmental stage they are capable of interpretation, judgement and modulation of their subjective experiences. Expression of the SIIPS1 signature may vary according to the functional development of key brain regions such as the ventromedial prefrontal cortex and nucleus accumbens, and the limited ability of infants to engage these systems in processes such as descending modulation^5^ and self-regulation^24^. A lack of concordance of infant brain activity with the SIIPS1 was primarily driven by divergent responses in ‘anti-pain’ brain regions, which occurs when a brain region that is normally de-activated with increasing pain intensity is instead activated, for example in patients with fibromyalgia^19^. Divergence of responses within the ‘anti-pain’ component of the signature, including regions such as the cuneus, hippocampus, and areas of the precentral gyrus, was also seen in the NPS, and was the most striking axis of differentiation between infants and adults. It is possible that immaturity of the haemodynamic response in infants produces altered BOLD responses in these areas. However, while differences in BOLD mechanisms have been observed in infant rats^25^, we used a haemodynamic response function derived from somatosensory responses in newborn human infants, where differences compared to adults appear more subtle^18^. Alternatively, the divergence in ‘anti-pain’ responses may be underpinned by true differences in noxious-evoked neural activity, where, for example, association and attentional networks are known to be immature in infants^26^. The positive evoked BOLD response within ‘anti-pain’ brain regions in infants could also potentially be due to exuberant connections of the infant brain^26^, coupled with a potentially underdeveloped GABAergic inhibitory system^27^. Some differences between infant and adult EEG responses to nociceptive stimuli have been reported, including delays in event-related potentials (ERPs), and a late ERP component in infants corresponding to an increase in delta band energy^20^. However, these explanations are speculative considering the exact mechanisms that underpin differences in the evoked haemodynamic activity in the infant and adult brain remain unclear.

Although caution is advised when translating and interpreting adult signatures in the immature brain^28^, there is considerable evidence to support the validity of this approach. Core structural features of the mature adult brain are developed in term-aged infants^29^ and functional responses^30^, connectivity patterns, and resting state activity^17,31^ recorded in infants exhibit similar features to adult brain activity. Furthermore, strong neural activity overlap between adult and infant sensory-evoked and noxious-evoked brain activity patterns have been reported across a variety of brain imaging modalities^6,20^. Despite the advantages of fMRI, it is important to appreciate that a single modality, cannot capture the full complexity of processes underlying a conscious experience. Therefore, we continue to advocate a multimodal measurement approach to provide the best proxy of infant pain^32^. Infant behavioural pain scores have been extensively studied, and brain-derived EEG signatures have been linked to infant nociception^20,21^. Linking these approaches to validated fMRI signatures in future studies will enhance our confidence in these methods.

The translational application of validated adult brain signatures to the immature brain holds great potential. This work provides the proof of concept for an approach to building more specific infant brain-derived biomarkers of pain. Given the clear structural and functional similarities in the organisation of the term infant and adult brain, projecting brain signatures onto the infant that have been refined based on adult verbal pain descriptors improves the inferences that can be made about infant pain. A particularly interesting application of this work would be to compare the representation of these signatures in prematurely-born infants who have reached term and term-born infants to assess how early life experiences associated with premature birth specifically impact pain experience. There is also value in applying the signatures to older infants and in children who are capable of verbally reporting their pain experiences, to investigate the developmental trajectory of the brain signatures. These signatures could be further optimised for infants by testing and validating the signatures in experimental paradigms in adults that are designed to replicate the reduced contextual understanding experienced by infants^33^.

This study expands the use of brain signatures to infer and deconstruct the experience of pain in a non-verbal patient population. Identifying the best proxy to quantify pain experience in infants is critical to the management of pain in neonatal care and the mitigation of potential long-term and short-term effects of pain in early life. This work provides a framework to better understand the early development of human sensory and emotional experience and provides a potential novel approach to assess the impact of analgesic interventions in infancy.

## Author Contributions

E. Duff, T. D. Wager and R. Slater designed the study. F Moultrie, S Goksan, M van der Vaart collected the data. E. Duff, A. Abos, S Fitzgibbon, L. Baxter and T. D. Wager analysed the data. E. Duff, F. Moultrie and R. Slater wrote the manuscript. All authors contributed to data interpretation and critically reviewing the manuscript.

## Supplementary Material (Methods)

### MRI acquisition

For adults, BOLD images were acquired using a T2*-weighted EPI acquisition (sequence parameters: TR/TE = 3280/40ms; flip angle = 90°; FOV = 192×192 mm; imaging matrix 64×64; resolution 3×3×3 mm; slices = 50, collected in descending order; average total volumes = 142). For Infant Cohort A, the same fMRI sequence was used with a reduction in slices (33 slices) and an infant-optimised TR/TE of 2500/40ms ^1^. For Infant Cohort B: BOLD images were acquired using a T2* BOLD-weighted, GRE acquisition with EPI readout, 70° flip angle, TE= 50 ms^1^, TR= 1,300 ms, mean TA= 6 mins (approx.), multiband 4^2,3^, 90 × 90 in-plane matrix size, 56 slices, 2 mm isotropic voxels. All studies used a 32-channel head coil.

### MRI Pre-processing

For the infant data boundary-based-registration (BBR) was used to register functional data to high-resolution structural data. Dynamic motion-by-susceptibility distortions were corrected using FSL’s EDDY^4,5^, and ICA-based denoising was used to remove additional artefactual signals^6,7^. Additional regressors modelling periods of excessive motion-related signal as identified by DVARS^8^ were also included.

Parameter estimate maps for individual subject and intensity levels were transformed into adult MNI space via a series of transformations generated using ANTs’s SyN (Advanced Normalization Tools Symmetric image Normalization method)^9^. Multi-modal non-linear registration was first used to register structural images T2 to the age-matched T2 template^10^. This was combined with the appropriate atlas week-to-week nonlinear transforms to transform to 44 week template, and then to MNI standard space with the NIHPD2 lifespan atlases (ages birth to 21 years) used as intermediate transform targets^11,12^. For adults, FNIRT nonlinear registration was used to transform data to MNI space^13^.

### Signature analysis

The SIIPS1 signature was trained using data from which the NPS had been removed via regression. We performed the same procedure here. The CANLAB toolbox provided a pipeline for processing, quality control and visualisation of parameter maps for the signature-based analysis of our group fMRI data. This included the extraction, assessment, regression and of white matter and CSF components, the calculation of signature similarity measures, estimation of contrasts and SVM classifier maps, and NPS subregion analysis. We adapted CANLAB code to include regression of the NPS prior to SIIPS1, and analysis of SIIPS1 subregions and single trial variability https://github.com/canlab.

## Notes

### Competing Interest Statement

The authors have declared no competing interest.

### Summary of Updates

Text has been reorganised and clarified.

